# Investigating the effects of pre-stimulus cortical synchrony on behavior

**DOI:** 10.1101/2020.02.27.967463

**Authors:** Mats W.J. van Es, Joachim Gross, Jan-Mathijs Schoffelen

## Abstract

Rhythmic brain activity may provide a functional mechanism that facilitates dynamic interareal interactions and thereby give rise to complex behavior. It has been shown that low and high frequency oscillations propagate in opposite directions, but interactions between brain areas in various frequency bands are poorly understood. We investigated local and long-range synchrony in a brain-wide network and their relation to behavior, while human subjects executed a variant of the Simon task during MEG recording. We hypothesized that the behavioral difference for stimulus-response congruent (C) and incongruent (IC) trials is caused by differences in cortical synchrony, and that the relative behavioral benefit for trials following instances with the same stimulus-response contingency (i.e. the Gratton effect) is caused by contingency-induced changes in the state of the network. This would be achieved by temporarily upregulating the connectivity strength between behaviorally relevant network nodes. We identified regions-of-interest that differed in local synchrony during the response phase of the Simon task. Within this network, spectral power in none of the nodes in either of the studied frequencies was significantly different in the pre-cue window of the subsequent trial. Nor was there a significant difference in coherence between the task-relevant nodes that could explain the superior performance after compatible consecutive trials.

## 1. Introduction

The brain is a complex organ that allows an organism to respond dynamically to an infinite set of inputs. In this context, the brain is considered to consist of a network of functionally specific units that dynamically interact to give rise to perception, cognition, and behavior. Rhythmic brain activity, a ubiquitous feature of neuronal activity, may provide a functional mechanism that facilitate these dynamic interareal interactions. Experimentally, rhythmic activity has been implicated in many cognitive tasks, where different frequency bands facilitate specific functions. It is thought that this characteristic is one of the main building blocks of brain signaling, but the interactions between different cortical areas and frequency bands is poorly understood.

In recent years, several studies found evidence for opposing propagation directions for oscillations in the high and low frequency bands. Gamma oscillations (30-90 Hz) have been found to propagate in a feedforward direction, transmitting sensory signals. Lower frequencies, mostly in alpha (8-12 Hz) and beta (12-30 Hz) bands, propagate in the feedback direction and mediate feedforward signaling. For example, van Kerkoerle and colleagues (2014) found that microstimulation in monkey area V1 elicited gamma oscillations in area V4, a higher level visual area. Moreover, stimulation in area V4 induced alpha oscillations in area V1. Studies using frequency-specific measures of directed influences found similar results (Bastos et al., 2015; Michalareas et al., 2016; Richter et al., 2017). In addition, gamma oscillations are thought to reflect high neuronal excitability (Fries, 2015; Schroeder and Lakatos, 2009), and brain regions exhibiting high gamma activity have been found to positively affect stimulus processing, and with that, behavioral performance (van Es and Schoffelen, 2019). On the other hand, alpha oscillations have been associated with inhibition of task-irrelevant regions (Jensen and Mazaheri, 2010), are especially apparent in (visual) spatial attention tasks (Bauer et al., 2014; Doesburg et al., 2016; Lobier et al., 2018), and have been shown to modulate the amplitude of gamma oscillations (Roux et al., 2013; Spaak et al., 2012). Beta oscillations are thought to have similar top-down functions, mostly in the sensorimotor domain (Engel and Fries, 2010). Taken together, increased gamma activity during an experiment is often interpreted as increased active processing of the task, while alpha and beta mediate this activity, and suppress irrelevant processing.

Yet, the understanding of functional interactions between brain areas and the role played by specific frequencies is incomplete. Many informative insights have been obtained from animal studies (Bastos et al., 2015; Richter et al., 2017; van Kerkoerle et al., 2014), which may not generalize to humans. Supporting evidence from human studies is limited, and confined to specific cognitive tasks (Popov et al., 2018; Schoffelen et al., 2017). Moreover, the human literature is often obtained from patient populations (Canolty et al., 2006). In the current study, we focus on changes in local and long-range synchrony in a brain-wide network involving the visuo-motor and attentional domains. We investigated whether these neuronal measures relate to subsequent behavior. Our intention was to investigate whether the state of a task-relevant brain network, both in terms of local oscillatory activity and of long-range synchronization, is directly related to behavioral efficiency, as indicated by the response speed during a variant of the Simon task. Subjects were required to make a speeded response with either the left or right hand after the onset of a visual response cue indicating which response to make. The classical Simon effect is characterized by differences in behavioral performance between stimulus-response congruent (i.e. instructive stimulus and instructed response are on the same hand side), and stimulus-response incongruent trials (Simon and Rudell, 1967). Interestingly, the Simon task effect size is susceptible to sequential dependencies, which is known as the congruency sequence effect or Gratton effect (Gratton et al., 1992). In short, the Gratton effect reflects an interaction of the behavioral benefit of stimulus-response congruency of a given trial, based on the stimulus-response congruency of the directly preceding trial (Gratton et al., 1992; Stürmer et al., 2002). We hypothesize that the neural basis for the Gratton effect may manifest itself in the period after the completion of the previous trial, before the presentation of the next stimulus. Specifically, the communication-through-coherence (CTC) hypothesis states that efficient neuronal communication relies on synchronization (Fries, 2015) in a network of behaviorally relevant brain areas. Faster responses might not only result from more efficient local information processing, but also from more efficient information transfer between brain areas. Therefore, optimal performance might be achieved when connections between task-relevant areas are more strongly synchronized at the moment at which new stimulus information becomes available. We hypothesize that the Gratton effect might be in part explained by modulations in interareal synchronization induced by the stimulus-response contingency of the previous trial. Specifically, a stimulus-response congruent trial would temporarily bias the strength of synchronization in intrahemispheric visual to motor connections, thus facilitating faster responses upon subsequent congruent trials. Likewise, a stimulus-response incongruent trial would temporarily bias the strength of interhemispheric visual to motor connections, thus facilitating less slow responses upon subsequent incongruent trials.

## 2. Methods

### 2.1 Subjects

19 healthy volunteers participated in this study, of which 6 male and 13 female. Their age range was 18-35 (mean ± SD: 22 ± 4.5). All subjects had normal or corrected-to-normal vision, and all gave written informed consent according to the declaration of Helsinki. This study was approved by the local ethics committee (University of Glasgow, Faculty of Information and Mathematical Sciences) and conform to the Declaration of Helsinki.

### 2.2 Experimental design

#### 2.2.1 Stimuli

The experiment was performed using Presentation^®^ software (Neurobehavioral Systems, Inc., Berkeley, CA, www.neurobs.com; RRID:SCR_002521). Throughout the experiment, a white, Gaussian blurred fixation dot was presented in the center of the screen against a black background (figure 1). After a 1.5 s baseline period, two white dots were presented bilaterally at equidistance from the fixation dot for 1-1.5 s, which functioned as warning cues. The response cues could either be both full circles, or one full and one half circle, in which case the half circle could surround the left or right part of either warning cue. Four of these response cue – warning cue combinations were presented before returning to the black background with only the fixation dot.

**Figure 1.**
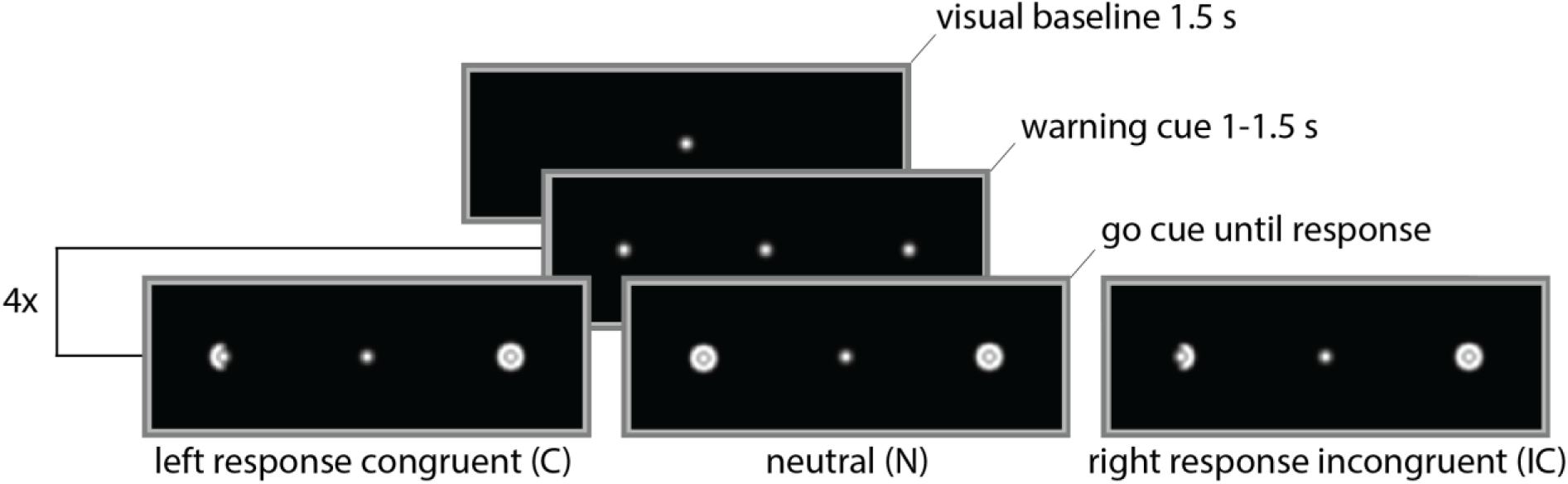
Task time line. A baseline period (1.5 s) was presented preceding every four trials, consisting of a fixation dot on a black background. The warning cue (1-1.5 s, jittered) preceded one of five different response cues. Displayed here: left response congruent (C), neutral (N) and right response incongruent (IC). Other response cues were left response incongruent or right response congruent. The response cue remained on the screen until a response was made.

#### 2.2.2 Experimental equipment

Brain signals were recorded with a 248 magnetometer 4D-Neuroimaging MAGNES 3600 WH MEG system, sampling at 1017 Hz, with online 0.1 Hz high-pass filtering. The MEG system was situated in a magnetically shielded room. Head position was assessed via five coils attached to the subject’s head. In order to co-register the MEG and MRI data, the scalp surface was digitized (FASTRAK^®^, Polhemus Inc., VT, USA), together with the five head position coils and three anatomical landmarks (nasion, left, and right pre-auricular points). Stimuli were presented through a DLP projector (PT-D7700E-K, Panasonic) situated outside the magnetically shielded room, onto a projection screen via a mirror inside the room. Additionally, anatomical T1-weighted MRI scans of the brain were acquired with a 3T Siemens MRI system (Siemens, Erlangen, Germany).

#### 2.2.3 Procedure

Subjects were instructed to fixate their gaze on the fixation dot in the center of the screen (see figure 1). Every four trials were preceded by a baseline window, in which only the fixation dot was present. Then, two warning cues appeared on either hemifield for 1-1.5 seconds, after which two response cues were presented, one in each visual hemifield. This is a modification of the classical Simon-task paradigm, in which just a single response cue is presented in one of the hemifields. Our stimulus presentation scheme required active processing in both visual cortical hemispheres, and the stimuli were designed to be very similar to one another, to ensure low-level visual processing to be as similar as possible across conditions. The response cues instructed the subject how to respond: if both response cues were full Gaussian blurred circles, no response was required (neutral condition, N). Alternatively, one of the response cues consisted of a full circle, while the other was a half circle. If the half circle was present on the left (right) side of the warning cue, the subject had to respond with a left (right) hand button press. Trials in which the response hand was on the same side as the warning cue are considered congruent (C); trials in which there is a mismatch between the informative hemifield and the response hand are incongruent (IC). All five conditions (neutral, and left/right hand congruent/incongruent) had 168 trials, making a total of 840 trials, presented in random order. In addition, a 4.5-minute empty room recording was acquired.

### 2.3 Data analysis

All data were analyzed in MATLAB 2017b (Mathworks, RRID SCR_001622) using FieldTrip Toolbox (Oostenveld et al., 2011; RRID SCR_004849) and custom written code. All results were visualized using Matlab or FieldTrip plotting functions, the RainCloud plots tool (Allen et al., 2019), and the circularGraph tool (Kassebaum, 2019). All experimental data and analysis scripts can be accessed from the Donders Repository (http://hdl.handle.net/11633/aabghkjl).

#### 2.3.1 MEG preprocessing

Power line interference at 50 Hz (with 0.2 Hz bandwidth) and its harmonics were removed from the data with a discrete Fourier transform (DFT) filter. To allow for the monitoring of the head position of the subject, the head localization coils were continuously activated throughout the measurement with a strong sinusoidal signal at 160 Hz. This signal was recorded along with the MEG, and regressed out of the MEG signals. Segments of the data containing SQUID jump artifacts or eye blink artifacts were identified and removed from the data. Ambient noise was reduced by regressing out the signals recorded by a set of reference sensors, located in the top of the MEG dewar. Lastly, the data were resampled to 300 Hz.

Additionally, the 4.5-minute empty room MEG recording was preprocessed as follows. The data were cut up in 2-second snippets with 50 % overlap, and demeaned. Excessively noisy snippets were removed from the data based on the variance. Then, the data were resampled to 300 Hz.

#### 2.3.2 MRI preprocessing

MRI data were co-registered to the MEG coordinate system using the co-registration information from the digitized head surface. For each subject, a 3D source model with an approximate 4 mm resolution was created with SPM8 (Penny et al., 2011), leading to 37,173 dipole locations inside the brain compartment. Additionally, volume conduction models were created using a single shell model of the inner surface of the skull (Nolte, 2003).

#### 2.3.3 Data stratification

Before the spectral analysis of the post-cue window, the data were stratified for data length and reaction times. Since power is positively biased by the number of samples, conditions with overall more samples (i.e. containing trials with larger reaction times) are confounded. First, the number of trials were equalized across conditions, and the remaining trials were matched for number of samples, cutting off the end of the trials with more samples. This operation ensured the same amount of data in each condition per contrast. However, this caused the data to be potentially confounded by latency: trials with higher RTs could have the same spectral characteristics as low RT trials, but later in time. Untimely cutting off these trials could abolish these spectral effects and confound the contrast. Therefore, trials from congruent and incongruent trials were removed at random until their distributions of RTs were equal, ensuring an overall comparable timing over conditions.

#### 2.3.4 Spectral analysis

Spectral analysis was performed in six a priori defined frequency bands: theta, alpha, beta and three frequency bands in the gamma range (see table 1). Spectral estimates of center frequencies were averaged within frequency bands. A 500 ms time window was selected (200-700 ms post-response-cue onset or until a response was made, or 500-0 ms pre-response-cue onset) and the data were detrended and zero padded to 2 s. Spectral analysis was performed at the particular frequency bins with the respective smoothing parameter (table 1), using Fast Fourier Transform (FFT) and DPSS multitapers in order to get the desired spectral smoothing. Because of the short time window in the post-cue time interval, theta and alpha power in this window could not be estimated with the desired smoothing (data segments were mostly below 500 ms), and was therefore estimated with 4 Hz smoothing in all trials. For the analysis in the pre-cue window, trials that were preceded by a baseline were removed from the data, because the baseline interval potentially dilutes potential congruency sequence effects.

**Table 1.** Definition of frequency bands, with the corresponding frequency bins used for estimation, and desired spectral smoothing. For the gamma bands, power was estimated at two center frequencies, and later averaged.

| Frequency band | Frequency range (Hz) | Center frequencies (Hz) | Smoothing (Hz) |
| --- | --- | --- | --- |
| Theta | 4-8 | 6 | 2 (or 4 for post-response cue) |
| Alpha | 8-12 | 10 | 2 (or 4 for post-response-cue) |
| Beta | 14-30 | 22 | 8 |
| low Gamma | 30-50 | 38, 42 | 8, 8 |
| mid-range gamma | 50-70 | 58, 62 | 8, 8 |
| high Gamma | 70-90 | 78, 82 | 8, 8 |

The empty room data were processed similarly, and were used for spatial pre-whitening of the frequency data. This reduces the influence of background interference in source reconstruction (Sekihara et al., 2006).

#### 2.3.5 Source reconstruction

Spectral power at the source level was estimated with dynamic imaging of coherence sources (DICS) beamforming. Pre-whitened frequency data was concatenated over conditions, and in case the window of interest was after response-cue onset, concatenated over pre- and post-cue windows. In case the window of interest was the pre-response-cue window, only the pre-cue data were concatenated. A common spatial filter was estimated for every source location, using a beamformer with fixed dipole orientation, and a regularization parameter of 100% of the mean sensor level spectral power. The common spatial filter was subsequently used to estimate power for each condition separately.

#### 2.3.6 Definition of ROIs

After statistical evaluation of the power difference between congruent and incongruent trials in the post-cue window, regions of interest (ROI, see table 2) were defined based on this effect. These regions were defined in order to reduce the search space in later analyses (power and coherence effects in the pre-cue window). ROIs were defined separately for each frequency band that showed a post-cue power effect, even if multiple frequency bands showed an effect in the same brain area. Presumably, these frequency-specific ROIs within the same brain area increased sensitivity for further analyses.

**Table 2.** Locations of regions of interest per frequency. The MNI coordinates are given for trials that required a left hand response; for trials that required right hand response, these coordinates were mirrored in the sagittal plane.

| Frequency band | Region | Hemisphere (relative to response hand) | MNI coordinates (mm) |
| --- | --- | --- | --- |
| Theta | frontal | midline | 2 28 36 |
| Alpha | occipital | ipsi | -22 -100 -4 |
|  |  | contra | 18 -92 4 |
| Beta | occipital | ipsi | -38 -92 8 |
|  | parietal | ipsi | -14 -92 48 |
|  | sensorimotor | ipsi | -42 -40 60 |
|  |  | contra | 50 -24 56 |
| mid-range Gamma | occipital | ipsi | -34 -92 4 |
|  |  | contra | 30 -100 4 |
|  | parietal | contra | 30 -88 44 |
|  | sensorimotor | ipsi | -42 -52 68 |
|  |  | contra | 62 -32 44 |
| high Gamma | occipital | ipsi | -22 -100 12 |
|  |  | contra | 30 -92 40 |
|  | parietal | ipsi | -26 -76 64 |
|  |  | contra | 26 -60 80 |
|  | sensorimotor | contra | 42 -32 68 |

#### 2.3.7 Coherence

Coherence was computed only for the pre-cue data, between the previously defined ROIs, and separately for congruent and incongruent trials. First, the cross-spectral density (CSD) was estimated at the sensor level, based on the spectral analysis described before (see 2.3.3). The source level CSD for all ROI dipole pairs was computed by combining the dipoles’ common spatial filters with the channel level CSD. From this, coherence between ROIs could be computed. Coherence was computed for all combinations of ROIs in which in at least one of them a post-cue power effect was present (e.g. if two ROIs were defined based on the power effect in two gamma bands, coherence between them would only be estimated in those frequency bands). Additionally, connections between ROIs in the same brain area (e.g. between alpha ROI and gamma ROI, both in contralateral occipital cortex) were excluded.

#### 2.3.8 Single-trial analysis

In order to test whether pre-cue gamma power in the sensorimotor area correlated with reaction times, single-trial power was estimated on the locations with the highest positive/negative t-value in that area, based on the pre-cue whole brain power analysis. Spatial filters that were retrieved from source reconstruction (see 2.3.5) were multiplied with the Fourier spectrum in order to get single-trial power. These power values were averaged over the frequencies within the low gamma band (38 and 42 Hz), correlated with reaction times, and subjected to a dependent samples T-test on the group level, contrasting the correlations to zero.

### 2.4 Statistical analysis

#### 2.4.1 Behavior

The effect of stimulus response congruency on the current trial’s reaction time (RT) was estimated with a two-tailed T-test. The effect size was defined as the behavioral advantage of congruent over incongruent trials in terms of seconds and percentages.

The effect of the congruency of the previous trial on the reaction time of the current trial was assessed with a two-way repeated measures ANOVA with two factors: congruency on the previous, and on the current trial. Congruency on the previous trial had three levels (congruent, incongruent, and neutral). Congruency on the current trial had only two levels (congruent and incongruent), since neutral trials did not require a response. Subsequently, partial eta squared was calculated to function as the estimated effect size.

#### 2.4.2 Spectral Power

From the single-trial source level spectral power, robust estimates of the condition-specific means and pooled variances were computed. Specifically, we computed a trimmed mean across trials, using a percentage of 20% trimming, and a 20% winsorized variance (Wilcox, 2011). This procedure yields more robust estimates when the data are not normally distributed, or when the data in the to-be-contrasted conditions have unequal sample sizes or variance. Sample means were converted to Z-scores, by means of normalization with the pooled standard deviation, to account for overall signal difference across subjects. This procedure was repeated for each frequency, and done separately for left hand and right hand response trials. Left hand and right hand responses were pooled by flipping the hemispheres of right hand response trials over the sagittal plane, and averaging the normalized sample means. Next, first level condition estimates were averaged within frequency bands (see table 1).

The normalized sample means were subjected to a nonparametric permutation test with 10,000 permutations based on the dependent samples Yuen-Welch T-statistic. In this case, the subject with the most extreme value in either tail of the distribution of that source location was excluded before calculating the samples means and variances. This step was done to account for overall signal differences across subjects. For the post-cue contrast, permutation testing was combined with spatial clustering for family-wise error control (Maris and Oostenveld, 2007). Adjacent dipole locations with T-values corresponding to a nominal threshold of 0.05 were grouped into clusters and their T-values were summed. The null-hypothesis was rejected if the maximum cluster statistic in the observed data was in either tail of the permutation distribution (with the critical alpha (0.05) Bonferroni corrected for six frequencies).

For the pre-cue contrast, 17 tests were carried out, one for each ROI in a specific frequency (see table 2). These tests evaluated if there was a consistent difference in the pre-cue window between trials preceded by congruent versus incongruent trials. The critical alpha level was Bonferroni corrected for multiple comparisons. Additionally, all effect size estimates are based on absolute power. For the post-cue window, the estimated effect size was based on the average effect size within the cluster that most contributed to the significant effect.

#### 2.4.3 Coherence

Coherence in congruent and incongruent trials were contrasted using a dependent samples Yuen-Welch T-statistic (as described above), and subjected to nonparametric statistics with 20,000 permutations. This was done separately for all 215 possible combinations of ROIs in a specific frequency band, and the critical alpha level was Bonferroni corrected for the number of comparisons.

## 3. Results

All nineteen subjects understood the task and performed well during the task. The average performance rate over subjects was 97% (SD = 3.0%) and the mean reaction time on correct trials was 651 ms (SD = 86.0 ms). Of all trials, on average 32 trials (SD = 24.5) contained artifacts in the MEG data and were removed from subsequent analyses.

### 3.1 Subjects respond faster when the previous trial was of the same congruency

The task was designed in order to elicit two well-known psychological effects: the Simon effect and the Gratton effect. The Simon effect presents itself as a difference in reaction times or performance when the instruction stimulus and the instructed response are on the same side, compared to when they are on opposite sides (Simon and Rudell, 1967). The current task presented a response cue for either left or right hand response in either left or right hemifield (and a non-informative stimulus in the opposite hemifield), and is thus expected to show a similar effect. Subjects were on average 44.4 ms (SD = 47.1 ms) faster on congruent trials than on incongruent trials, t(18) = −4.12, p=0.00065 (see figure 2a), which confirms the effect.

**Figure 2.**
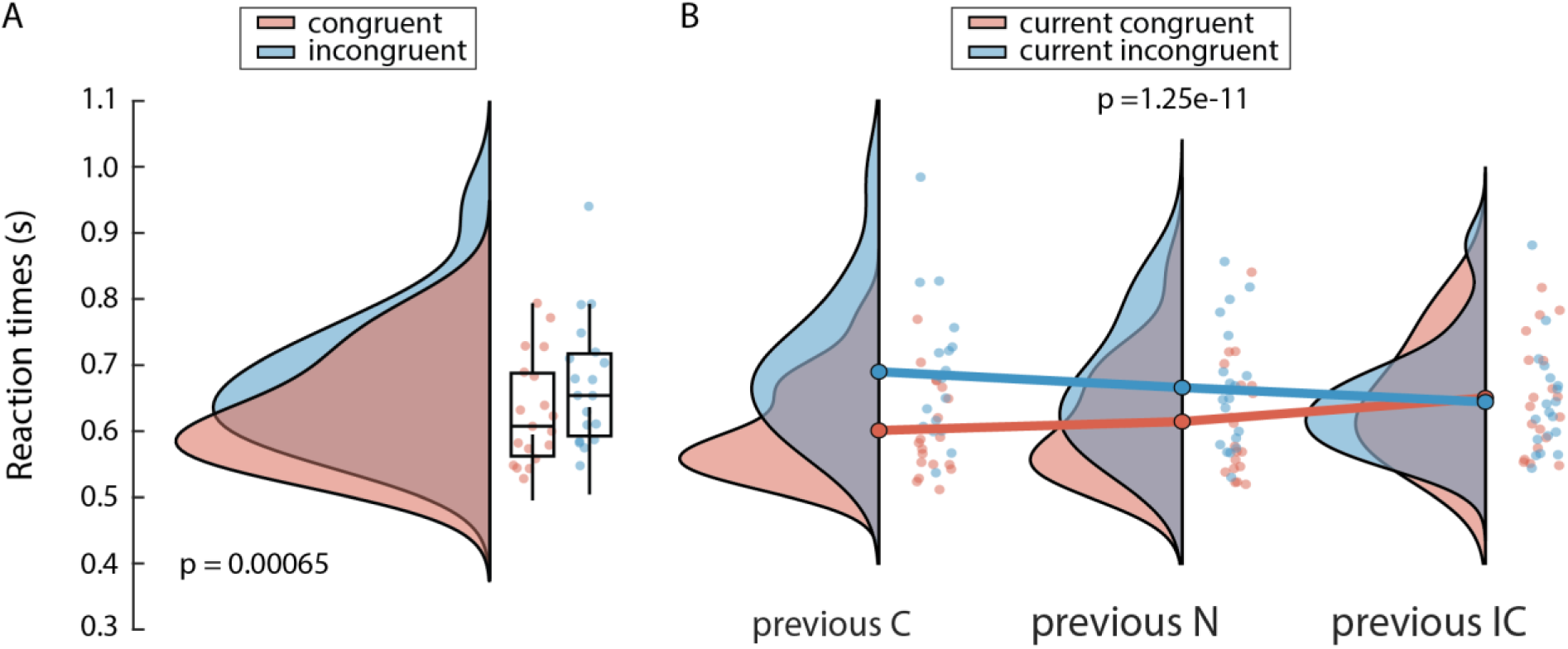
Reaction times differ based on stimulus-response congruency. A) Subjects respond faster on congruent than incongruent trials. B) Subjects respond faster when the previous and current trial are compatible. Graphs show probability density, box plots denote median, 1^st^ and 3^rd^ quartiles and 1.5 times the interquartile range (IQR). Dots correspond to data points of individual subjects.

The other expected effect is a congruency sequence effect, also known as the Gratton effect: the difference in behavioral performance for compatible versus incompatible sequential trials (i.e. of the same or different congruency), which is also present in Simon experiments (Notebaert et al., 2001). We tested whether the Gratton effect was induced by the task by testing the effect of stimulus congruency of the current trial on RTs, conditioned on the previous trial. Reaction times were lower when the current trial was preceded by a trial of the same congruency (F_(2,18)_ = 54.6, p = 1.25e-11, η^2^= 0.049; see figure 2b), e.g. subjects responded faster on congruent trials when the previous trial was also congruent, rather than incongruent, and vice versa. Interestingly, when a trial was preceded by a neutral trial, reaction times were in between reaction time of compatible and incompatible trials.

The confirmation of these two behavioral effects demonstrate that the experimental design caused the behavior as intended. In the rest of the analyses, we investigated the oscillatory neural correlates of these effects. First, the relevant frequency bands and network nodes for successful completion of the task were identified by investigating spectral power differences between congruent and incongruent trials, in the time window from response cue onset until the response. These resulted in a number of regions of interest (ROIs) that informed subsequent investigation of the Gratton effect. Spectral power in these ROIs, and coherence between them, were computed in the 500 ms time window before the onset of the response cue. These measures are assumed to reflect the state of the task relevant network, right before the informative response cue is presented. These neural states are hypothesized to affect the efficiency of task relevant processing in the subsequent task stage, thereby causing the observed behavioral differences.

### 3.2 The Simon effect is associated with differences in local neuronal synchrony in the visuo-motor network

We presumed that the behavioral advantage for congruent over incongruent trials can be (partially) explained by differences in neural synchrony related to response selection and initiation. This was tested by contrasting spectral power at the source level, between congruent and incongruent trials in six a priori defined frequency bands: theta (4-8 Hz), alpha (8-12 Hz), beta (14-30 Hz) and three frequency bands in the gamma range (30-50, 50-70 Hz, 70-90 Hz). For each subject we computed the difference in power between congruent and incongruent trials, separately for left and right handed responses. In order to increase statistical sensitivity, contrasts for left and right-handed responses were pooled after mirroring the spatial patterns of spectral power changes for right hand response trials in the sagittal plane, thus focusing on consistently lateralized responses. The resulting power differences (and all subsequent results) are therefore reported as ipsi- or contralateral to the response hand, and results are displayed as if all trials required a left hand response.

One potential confound for the contrast between congruent and incongruent trials is the overall higher reaction times for incongruent trials. Spectral power was estimated in the window from 200 ms to 700 ms after the response cue, or until a response was made. Power is not stationary within this window, and changes over time as a function of the neural process. Since incongruent trials were associated with longer RTs, the spectral changes likely arise later in time as well. If the analysis window captures different stages of the neural process depending on RT, this would lead to biased power estimates. Additionally, power in incongruent trials was estimated on larger time windows on average (because of longer RTs), leading to more accurate power estimates. In order to account for confound of reaction time, the analysis above was executed after stratifying conditions for data length and reaction times (see Methods 2.3.3).

Group statistical evaluation at the cluster level confirmed the presence of power differences between congruent and incongruent trials in three out of six frequency bands: beta (p = 0.036, nonparametric permutation test, corrected), mid-range gamma (p = 0.018, nonparametric permutation, corrected), and high gamma (p = 0.022, nonparametric permutation, corrected). The T-values of these contrasts are displayed in figure 3, and are qualitatively similar when the same analysis is done without stratifying the data (figure S1). The beta power effect is most clear in the sensorimotor cortex, where the effect is lateralized. It further extends to ipsilateral parietal and occipital cortex. In the cluster that lead to rejection of the null-hypothesis, power values were on average 4.3 % higher (SD = 2.6 %) in congruent trials compared to incongruent trials. The effects in the gamma range were also lateralized, generally in the opposite direction than the beta effect. Mid-range gamma was strongest in occipital cortex, with on average 3.5 % (SD = 2.4 %) higher power values in contralateral occipital cortex for congruent trials. The power effect in high gamma was present throughout the contralateral cortex, including sensorimotor, parietal and occipital cortex. On average, power values in the largest positive clusters were 3.6 % (SD = 3.3 %) higher in congruent trials.

**Figure 3.**
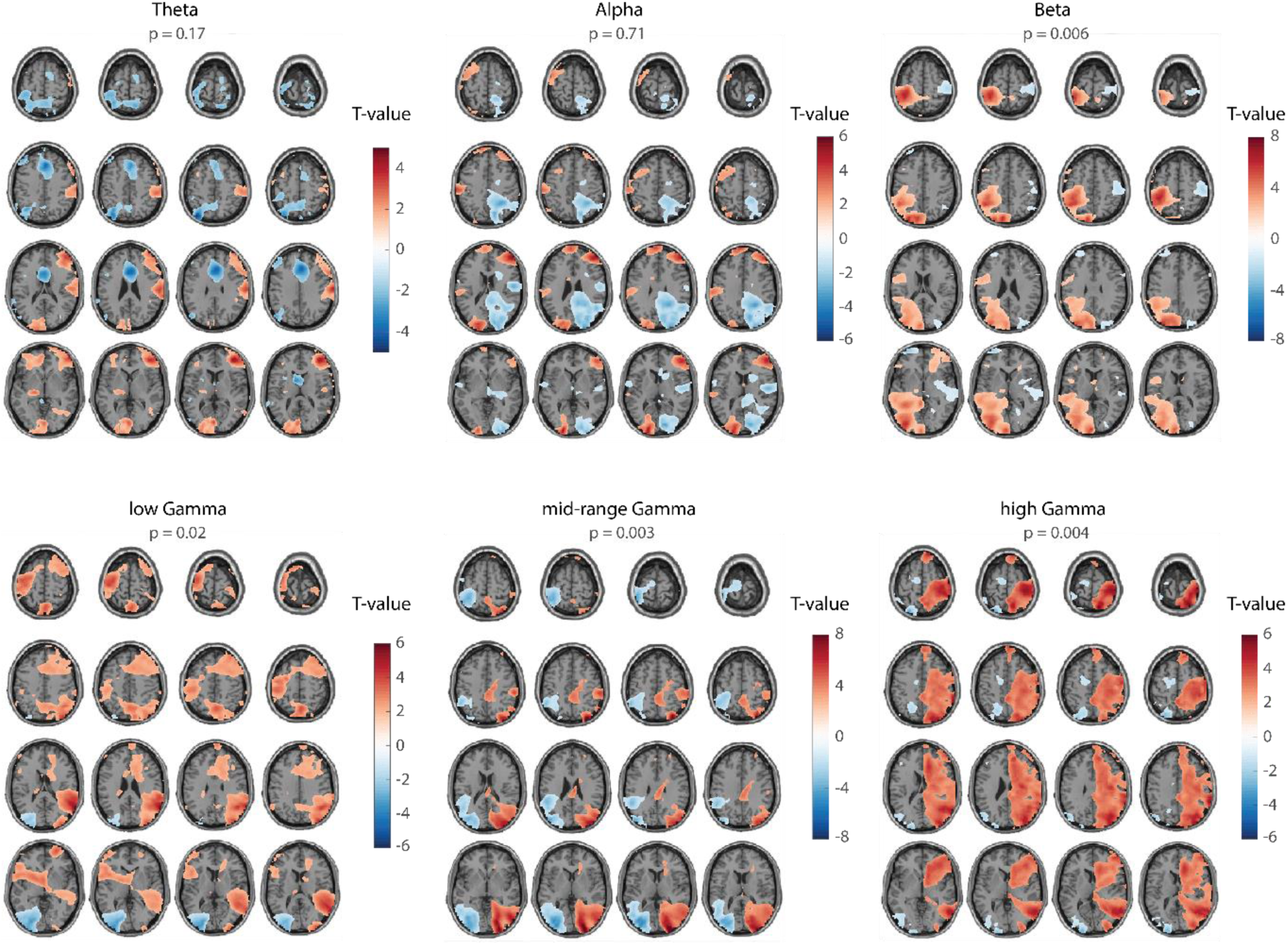
Spectral power differences in the post-cue window between congruent and incongruent trials, requiring a left hand response. The data were stratified on RT and data length before spectral analysis. Spectral differences are lateralized, and in opposite direction for low (alpha, beta) and high (gamma) frequency bands. Color values indicate T-values at the group level (values smaller than 30% of the maximum values are masked); p-values are not corrected for multiple frequencies.

In the frequency bands where the power difference was not significant (after correction); some interesting trends were visible. In the theta band (p = 0.17, nonparametric permutation test, uncorrected), congruent trials showed less power along the midline in frontal cortex. This might be related to processing conflicts (Cavanagh and Frank, 2014), which are especially present in incongruent trials. The trend in the alpha band (p = 0.71, nonparametric permutation test, uncorrected) showed lateralization in occipital cortex, notably in the opposite direction than the effects in gamma band, as is often seen in this brain area (Bauer et al., 2014; Jensen and Mazaheri, 2010). Lastly, the trend in the low gamma range (p = 0.02, nonparametric permutation test, uncorrected) was strongest in temporal cortex, but was present in small clusters throughout the cortex. It was the only frequency band that did not look qualitatively similar when the analysis was done without stratification of the data (figure S1), which make this result difficult to interpret.

Based on these observations, regions of interest (ROIs) were defined for further analyses. ROIs were considered in the three frequency bands in which a statistically significant post-cue power effect was present (beta, mid-range gamma, and high gamma). Additionally, ROIs in the theta and alpha bands were defined, despite the absence of a significant effect. In light of the literature (see Discussion), the trends present in these data point towards a possible involvement in the current task. Furthermore, possibly these trends did not reach significance because of the insensitivity of the current analysis in these frequency bands (see Methods 2.3.4 and Discussion). In total, seventeen ROIs were defined in specific frequency bands, as summarized in table 2.

### 3.3 Differences in oscillatory activity after congruent or incongruent trials only affect subsequent processing of identical trials

We presumed that the nodes identified in the post-cue window are generally important for task-relevant processing. Since there is a behavioral congruency sequence effect, it could be that activity in these task-relevant nodes, or connectivity between them, is adjusted according to the stimulus-contingency of a given trial, in order to rebalance the state of the network for optimal performance in the next trial. For example, after a congruent trial, functional connectivity within hemispheres would increase, at the cost of interhemispheric connectivity, essentially preparing the network for an efficient response in another congruent trial. The opposite is expected after an incongruent trial, where the task-relevant network would be prepared for another trial in which efficient interhemispheric information transfer is required. In order to test whether spectral power differed between conditions at trial onset (i.e. depending on the contingency of the preceding trial), power was estimated in the ROIs in the 500 ms time window before the onset of the next trial. Just as in the previous analysis, left and right hand response trials were pooled, and the congruency effect was tested statistically, conditioned on the previous trial. There was no consistent effect in either of the ROIs in any frequency (figure 4). In order to make sure that this was not a result of poorly defined ROIs, we also conducted a whole brain analysis, similar to the analysis in the post-cue window (figure 5). No statistically significant effect was present in any of the frequencies there either. There was a trend of higher theta power in ipsilateral parietal cortex and higher beta power in the mid-central area for incongruent trials in the whole brain analysis, but these were not studied in the ROI analysis. Another trend was visible in the alpha band in both the ROI and whole brain analyses, similar to the trend in the post cue window: more alpha power in ipsilateral occipital cortex after a congruent trial, and less power on the contralateral counterpart. The same trend was present in the mid-range gamma band, but in the opposite direction. These trends both point towards activation of the visual areas that received the informative stimulus in the previous trial, and inhibition of the areas that received the uninformative stimulus. Lastly, a trend was present in the low gamma band, with less power in ipsilateral parietal and sensorimotor cortices for congruent trials, and more power in the contralateral counterparts. This trend is not comparable to the contrast in the response window (figures 3 and S1), where the results were inconsistent for stratified and non-stratified data. However, it does suggest more excitation in the motor cortex that was responsible for the action in the previous trial, and less in the motor cortex that had to be repressed. This would indicate a behavioral benefit if the next trial required the *same* response, instead of a *compatible* response. Post-hoc behavioral analysis (figure S2) revealed that the Gratton effect is present if two sequential trials required the same response (F_(1,18)_ = 79.4, p = 5.11-8, η^2^ = 0.27), but not when they required a different response (F_(1,18)_ = 2.33, p = 0.14, η^2^ = 0.0025). It is therefore possible that the Gratton effect observed in our data is a consequence of trial repetition (i.e. two sequential trials are of the exact same type), rather than trial contingency. This opens up the possibility that the level of excitation in the motor cortex contralateral to the response hand, and/or the level of inhibition ipsilateral to the response hand determine the sequential trial effects in behavioral performance. However, there was no single-trial correlation of reaction times with power in either ipsi- or contralateral sensorimotor area (ipsilateral: t(18) = -0.13, p=0.90; contralateral: t(18) = 0.42, p=0.68).

**Figure 4.**
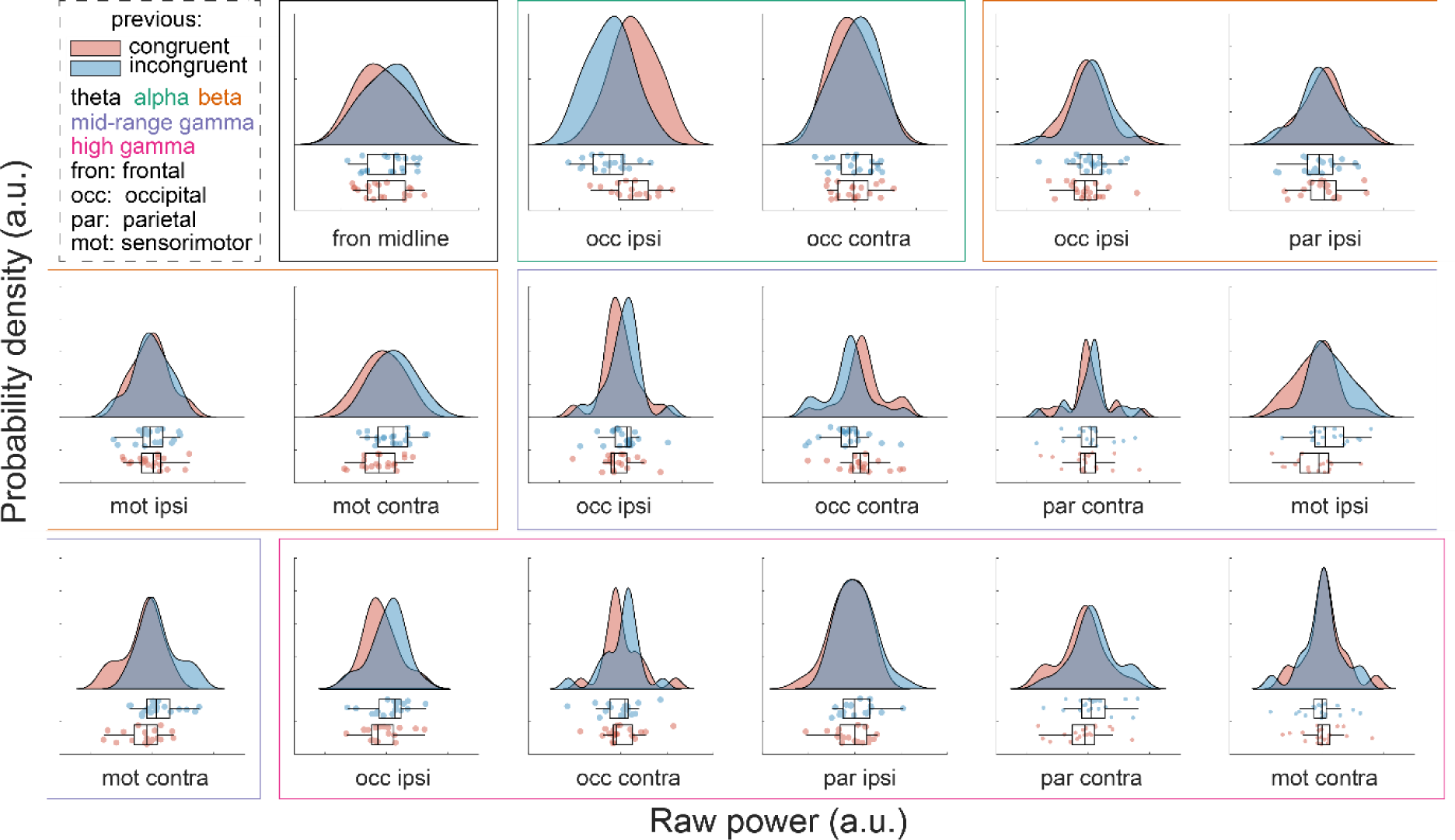
Contrast of power in congruent versus incongruent trials in the pre-cue window, in selected ROIs. There was no significant difference between congruent and incongruent trials in any of the ROIs. Graphs show probability density, box plots denote median, 1^st^ and 3^rd^ quartiles and 1.5 times the interquartile range (IQR). Dots correspond to data points of individual subjects.

**Figure 5.**
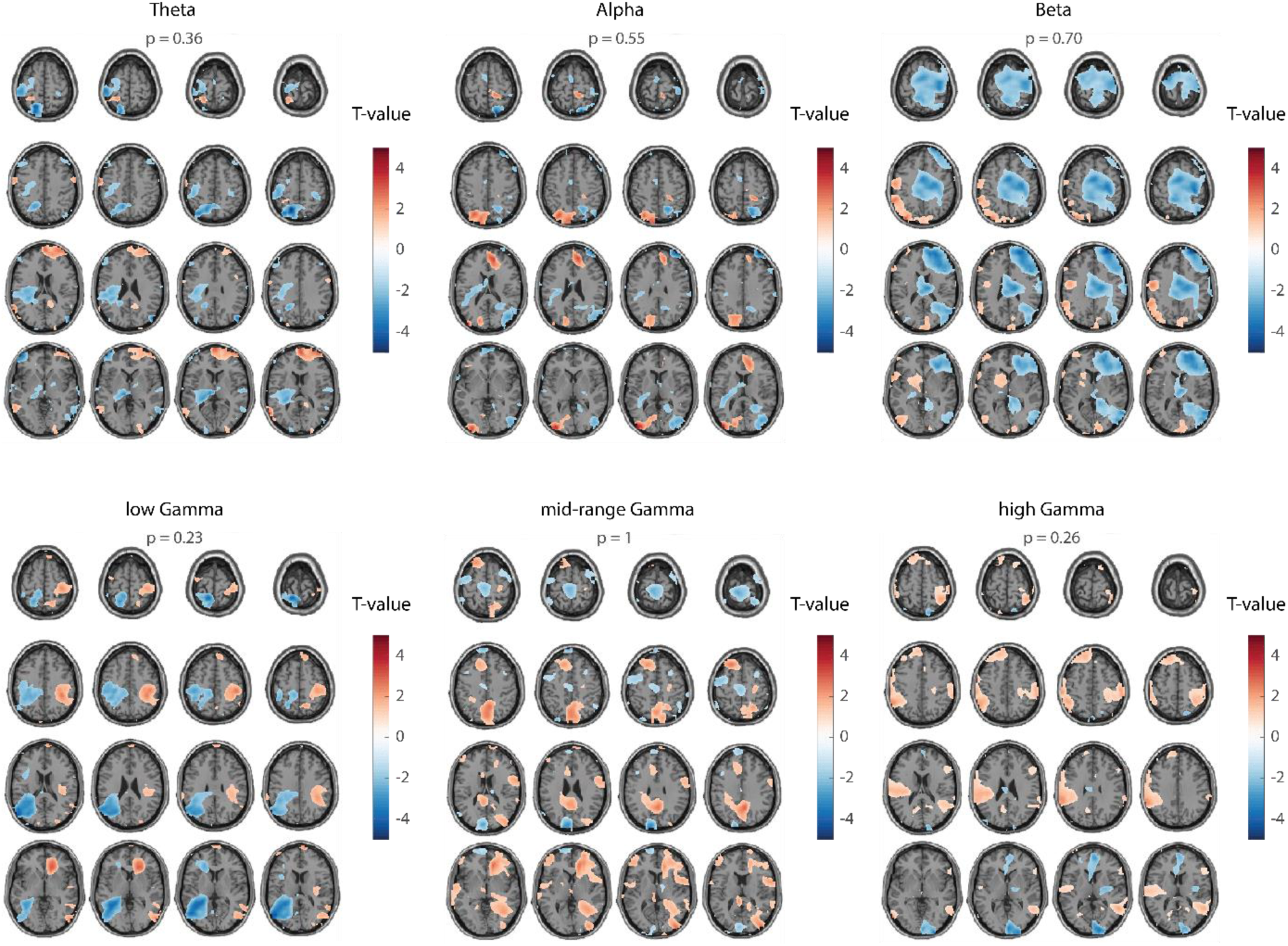
Whole brain spectral power differences in the pre-cue window between previous congruent and previous incongruent trials, requiring a left hand response. None of the frequency bands was significantly different. Color values indicate T-values at the group level (values smaller than 30% of the maximum values are masked); p-values are not corrected for multiple frequencies.

### 3.4 Differential interhemispheric connectivity might alter processing of subsequent trials

The absence of a statistically significant neuronal effect in the pre-cue window is at odds with the resulting behavior, where the response time is affected by the condition of the previous trial (i.e. Gratton effect). However, local neuronal synchrony does not represent the entire state of the network. Functional connectivity between network nodes, as indexed by phase synchrony, can potentially play a role in bringing about this behavioral effect, by facilitating efficient information transmission between areas. We hypothesized that high phase synchrony between task-relevant nodes leads to efficient information transfer. The phase synchrony in the current trial would still be present at the start of the next trial, and thus benefit trials that require the same network nodes to communicate (compatible), relative to two sequential trials where this is not true (incompatible). In particular, *intra*-hemispheric connections are thought to be stronger after congruent trials, whereas *inter*-hemispheric connections are thought to be stronger after incongruent trials.

In order to test this, we computed coherence between all pairs of ROIs in the pre-cue window. This was done for the five frequency bands used before, and separately for previously congruent and previously incongruent trials. Excluding connections that were not of interest (see Methods 2.3.7.), 215 connections were tested. None of them survived Bonferroni correction. Figure 6 depicts those connections with a p-value smaller than 0.05 before correction (see figure S2 for the remaining connections). Interestingly, most of these connections are interhemispheric and are stronger after incongruent trials. This could indicate that performance on incongruent trials increases when succeeding another incongruent trial, due to increased interhemispheric synchrony.

**Figure 6.**
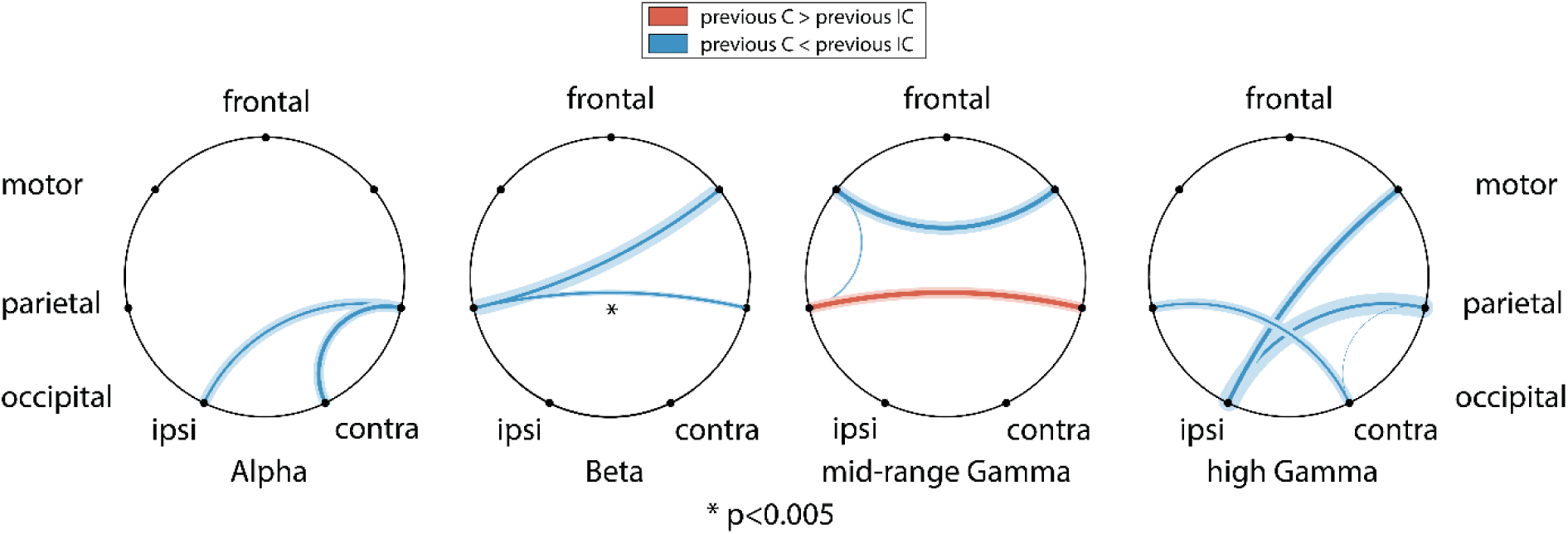
Coherence in the task-relevant network for all connections with p<0.05 (uncorrected). The relative difference in coherence for previous congruent versus previous incongruent trials in the pre-cue window. Solid lines indicate group average, with the thickness of the lines proportional to the ratio between coherence in congruent and incongruent trials; transparent lines indicate standard deviation. Red (blue) denotes larger (smaller) coherence for previous congruent versus incongruent. ipsi: ipsilateral to cued response hand; contra: contralateral to cued response hand.

## 4. Discussion

In this work, we investigated local oscillatory activity and long range functional interactions between brain areas in various frequency bands, while subjects were engaged in a Simon task. We hypothesized that the behavioral difference for stimulus-response congruent (C) and incongruent (IC) trials is caused by differences in cortical synchrony. Furthermore, we hypothesized that the relative behavioral benefit for trials following instances with the same stimulus-response contingency (i.e. the Gratton effect) is caused by contingency-induced changes in the state of the network, by temporarily upregulating the connectivity strength between behaviorally relevant network nodes. We identified regions-of-interest in the sensorimotor, visual, and attention-related areas that differed in local synchrony during the response phase of the Simon task. Within this network, using rigorous statistical procedures, spectral power in none of the nodes in either of the studied frequencies was significantly different in the pre-cue window of the subsequent trial. Nor was there a significant difference in coherence between the task-relevant nodes that could explain the superior performance after compatible consecutive trials.

The task we used in this experiment is not a standard Simon task. The original Simon task presented an auditory response instruction (left/right hand response) to either ear (Simon and Rudell, 1967). The current experiment was in the visual domain, and presented stimuli in both hemifields for each trial. Thus, the subject had to process both stimuli in order to identify the stimulus containing the response instruction, requiring engagement of bilateral early visual areas. Additionally, we included neutral trials, thus explicitly requiring the processing of both stimuli before a response decision could be made. These modifications to the original Simon task made the task more difficult, which is supported by longer RTs: the original study reported mean reaction times in the order of 400 ms, while on this task the average RT was 651 ms. The introduction of two visual stimuli with very similar low level features on each trial ensured that any differences between trials would unlikely be the result of trivial differences in the response of visual areas to low level stimulus characteristics.

We observed statistically significant power differences between C and IC trials in beta, mid-gamma and high gamma bands. The contrasts in theta and alpha band were not significant, despite the presence of clear trends that met our expectations. One reason why these two contrasts might not have survived statistical testing is the insensitivity of the analysis to low frequencies. Reaction times were still generally low, leading to short time windows on which power could be estimated. Consequently, spectral smoothing in low frequencies was higher than desired (4 Hz instead of 2 Hz). This lead to spectral leakage from frequencies outside of the frequency bands of interest that could have compromised sensitivity to the effects. Although no strong claims about the theta/alpha band could be made, we could interpret the trends in the data nonetheless.

The trend in the theta band was strongest in the mid-frontal region of the brain, where theta power was higher during IC trials than during C trials. Increases in frontal theta power are often observed during response conflict, likely reflecting the need for more cognitive control. In line with this, is has already been shown that power in a Simon task is higher for IC than C trials (Cohen and Ridderinkhof, 2013; Nigbur et al., 2011), and higher for IC trials following IC, rather than C trials (Pastötter et al., 2013).

Alpha power, likely reflecting inhibition, was reduced in the occipital cortex contralateral to the, and enhanced ipsilateral to the informative stimulus. This pattern is often observed in visual spatial attention tasks (Bauer et al., 2014; Jensen and Mazaheri, 2010; Thut et al., 2006). Whereas most spatial attention tasks cue the subject on which hemifield should be attended before the presentation of the relevant stimulus, in the current experiment the warning cue did not contain spatial information. All warning cues were also designed to be similar in their low-level visual features. This lateralization is probably top-down in nature and instantiates after at least partial processing of the stimuli, since it cannot be caused by attentional preparation and it is unlikely to be stimulus-induced.

Beta desynchronization is classically observed in sensorimotor cortex during movement execution (Neuper et al., 2006; Pfurtscheller, 1981; Pfurtscheller and Lopes da Silva, 1999), and the movement duration is shorter with larger desynchronization (Heinrichs-Graham and Wilson, 2016). Increases in beta synchronization have been observed if a prepared movement is terminated, for example in Go/No-go tasks (Alegre et al., 2004; Zhang et al., 2008). In the current study, C trials, relative to IC trials, showed lower beta power in the hemisphere contralateral to the cued response hand and higher beta power in the ipsilateral hemisphere. Faster reaction times for C compared to IC trials are in line with the larger beta desynchronization in the ipsilateral sensorimotor cortex. Additionally, higher beta power in the ipsilateral hemisphere suggests superior inhibition of the incorrect motor response in C trials.

Just like the occipital alpha lateralization, the occipital gamma lateralization (in the opposite direction compared to alpha) is unlikely to be stimulus-induced. Still, it is unlikely that this lateralization (or the one in alpha band) causes the behavioral difference in C and IC trials. Namely, the lateralization was present in both C and IC trials, but in opposite directions (data not shown). This is unsurprising because the informative stimulus is also present in opposite hemifields for these trial conditions. These results do therefore not point to a difference in visual processing between C and IC trials per se, but rather indicate that visual processing of the relevant stimulus is upregulated immediately after identification of its relevance. Similarly, processing of the irrelevant stimulus is immediately downregulated.

Further, gamma power was also lateralized in sensorimotor cortex, in the same direction as in occipital cortex, but mainly in the high gamma range (70-90 Hz). Sensorimotor gamma synchronization in this frequency range is reported to have a prokinetic role (Seeber et al., 2015), and gamma synchronization accompanies beta desynchronization upon sensorimotor activation (Crone, 1998). The power lateralization in sensorimotor cortex in opposite directions for beta and gamma therefore provides reinforcing evidence for enhanced sensorimotor activation of the correct motor response in C trials, and enhanced inhibition of the incorrect motor response.

Taken together, the rich pattern of spectral differences in the response window of the Simon task mainly point functional differences in the response preparation and execution that benefit C over IC trials. This is supported by stronger activation of the sensorimotor area responsible for the correct response, and stronger inhibition for sensorimotor area responsible for the incorrect response. These findings fit well with the widely accepted cognitive theory that attributes the Simon effect to differences in spatial coding in the response selection phase (Hommel, 2011).

In addition to the Simon effect, the task used here elicited a sequential dependency effect, or Gratton effect. Subjects responded faster on trials that were compatible with the condition of the previous trial. Therefore, the stimulus-response contingency induces a specific change in the neuronal state that influences the behavior in the subsequent trial. We hypothesized that this neuronal state is reflected in the local and/or long-range synchronization of the network. In short, areas and functional connections that were required for a trial of one condition are strengthened through synchronization, hereby affecting the initial network state at the start of the next trial and improving performance for a trial of the same condition. We found no significant differences in power at the start of a next trial. The most interesting trends were visible in the alpha and mid-range gamma bands, with a lateralization in occipital cortex in opposite directions, similar to what was visible in the response window. Additionally, there were trends of higher ipsilateral theta power in parietal cortex and higher midcentral beta power, after IC trials.

Even if a lateralization in excitation and inhibition in visual cortex persisted after the previous trial, it is unlikely that this could cause the difference in performance as seen in the Gratton effect. Particularly, if an imbalance of excitability in visual areas would benefit subsequent visual processing, it would probably do so only in trials where the response cue is presented in the same hemifield, rather than trials of the same contingency. The same can be said for the lateralization in sensorimotor areas in the low gamma band: if this reflects a difference in excitation, it would likely only benefit trials that require the same response. Both these trends are supported by behavior: the Gratton effect was only present in trials that required the same response, and trivially had the instructive stimulus in the same hemifield. For example, if a congruent, *left*-hand response trial was preceded by a congruent, *right-*hand response trial, performance was as good as when it was preceded by an *incongruent*, right-hand response trial. This puts the origin of the Gratton effect in question, which has occurred before in the literature (see Hommel, 2011). Many reports saw the Gratton effect disappear when accounting for full repetitions of trial condition (Puccioni and Vallesi, 2012; Schmidt and De Houwer, 2011). Thus, the Gratton might not instantiate because of contingency effects, but because of increased performance after stimulus-response repetitions. According to a cognitive theory by Hommel (1998), this might be due to stimulus-response binding in episodic memory. The trends in the current data hint in an alternative direction: differential excitability in visual and motor cortical areas affect subsequent processing. There was no correlation between pre-cue gamma power in sensorimotor areas and reaction times though, possibly because the analysis did not take baseline gamma power into account. The lack of significant effects make our alternative explanation for the Gratton effect merely speculation, and overall the results remain inconclusive. Possibly, cognitive control has alternative consequences on visuo-motor processing, as suggested by Pastötter et al. (2013). They specifically studied the oscillatory correlates of cognitive control in a response-priming task and found that reaction times were lower on IC trials with higher midfrontal theta power in the response period. Moreover, they found higher ipsilateral parietal theta power and midcentral beta power after IC trials, a trend that was also present in our data. Pastötter and colleagues found that in subjects where this effect was stronger, the weaker was the effect in midfrontal theta power during the response phase, although the mechanism behind this remains unknown. It is unclear why the pre-cue effects in theta and beta bands were not significant in the current study. Pastötter et al. used a different task, but it is not likely that their task induced a greater need for cognitive control: reaction times in the current experiment were generally higher, and had a greater effect of condition. If anything, this task was more difficult and if cognitive control is the key factor, it was higher in our task, reflected by reaction times. Possibly, the difference in sample size and/or analysis approach lead to the differences discussed here.

The indefinite conclusions drawn from the power results could have been cleared up by investigating the connectivity between network nodes. There was a tendency of higher beta band coherence between ipsilateral and contralateral parietal cortex after an IC versus a C trial. This could reflect increased communication for conflict resolution between network nodes important in motor planning. In general, most connections with a p < 0.05 were interhemispheric connections, and were stronger after IC trials, which we hypothesized. We did however, find no trends in the theta band or in any connections with the midfrontal cortex, despite theta’s implication in pre-cue power (Pastötter et al., 2013) and during trial processing of a Simon task (Cohen and Ridderinkhof, 2013). Even though some interesting trends were present in the coherence analyses, they too remain inconclusive. This be due to too conservative correction for multiple comparisons, and perhaps the pre-defined ROIs were sub-ideal. For example, the trends in pre-cue power in the ipsilateral parietal theta power and midcentral beta power were not further investigated in a coherence analysis. Most likely, the presumption that the key differences in post-trial processing between C and IC trials take place in the same network nodes that show the largest differences in trial processing was too constraining.

In conclusion, we found differences in the spectral neural response of congruent and incongruent trials in a Simon task, most notably related to motor planning and execution, and possibly cognitive control. The observed trends in beta and theta band power after trial offset are in line with the literature that predict a role for cognitive control that affects processing of the subsequent trial. On the other hand, the trends in gamma band point to a role in motor planning and execution, which would only benefit subsequent trials if they were of the same type, and is supported by behavior. It is therefore unclear to what extent control and response planning can account for congruency sequence effects, and whether this reflects conflict adaptation to begin with. Further research should therefore focus on the origins of the Gratton effect in order to constrain analyses into the underlying mechanisms of sequential dependency effects.

## Acknowledgements

This work was supported by The Netherlands Organisation for Scientific Research (NWO Vidi: 864.14.011), the Wellcome Trust (Senior Investigator Grant: 098433), University Hospital Wuerzburg (IZKF Gro3/001/19), and the German Research Foundation (DFG GR 2024/5-1). The sponsors had no role in the collection, analysis and interpretation of data; in the writing of the report; and in the decision to submit the article for publication.

## 6. Supplemental information

**Figure S1.**
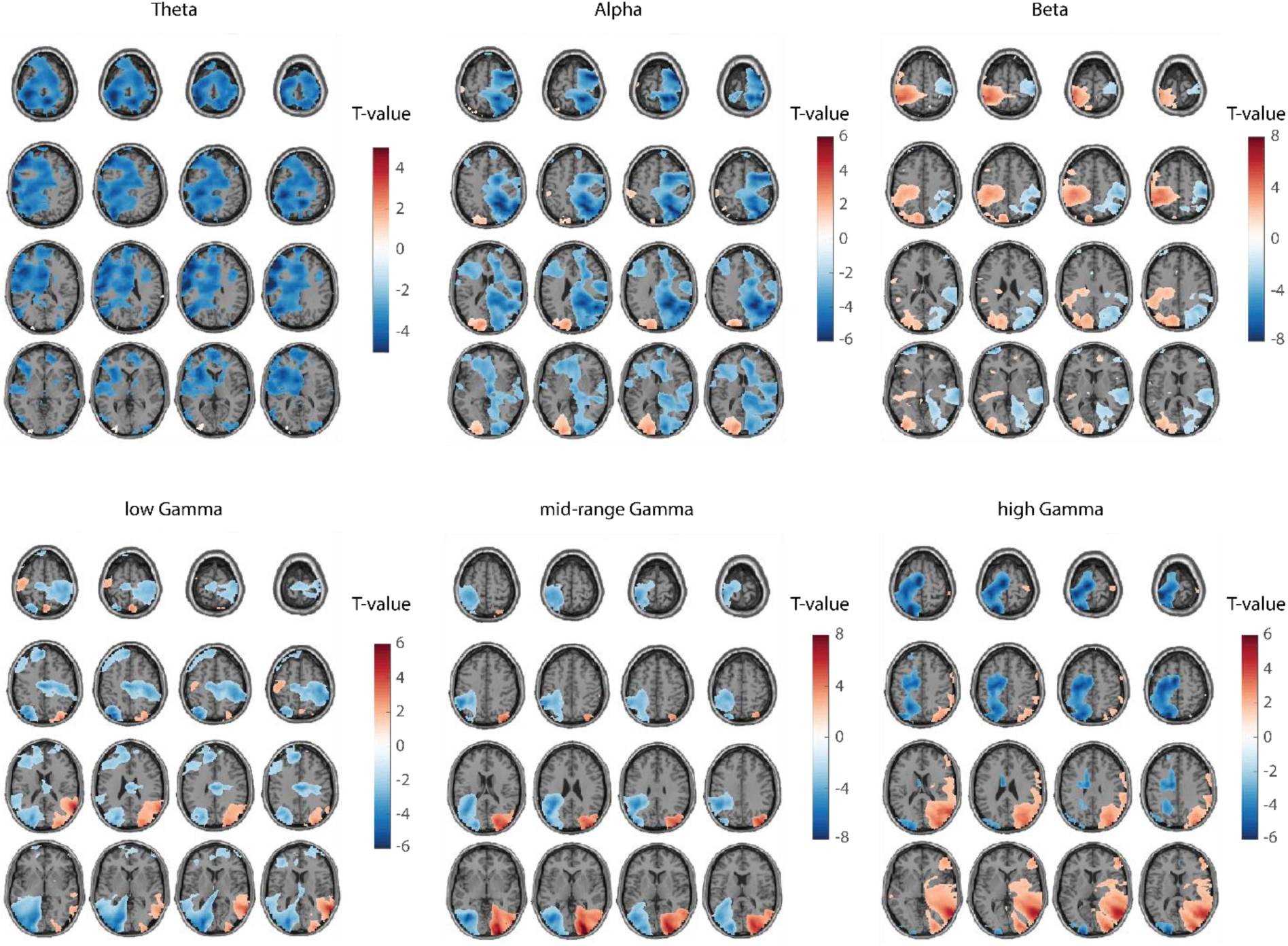
Spectral power differences in the post-cue window between congruent and incongruent trials, requiring a left hand response. Spectral differences are lateralized, and in opposite direction for low (alpha, beta) and high (gamma) frequency bands. Color values indicate T-values at the group level (values smaller than 30% of the maximum values are masked); p-values are not corrected for multiple frequencies.

**Figure S2.**
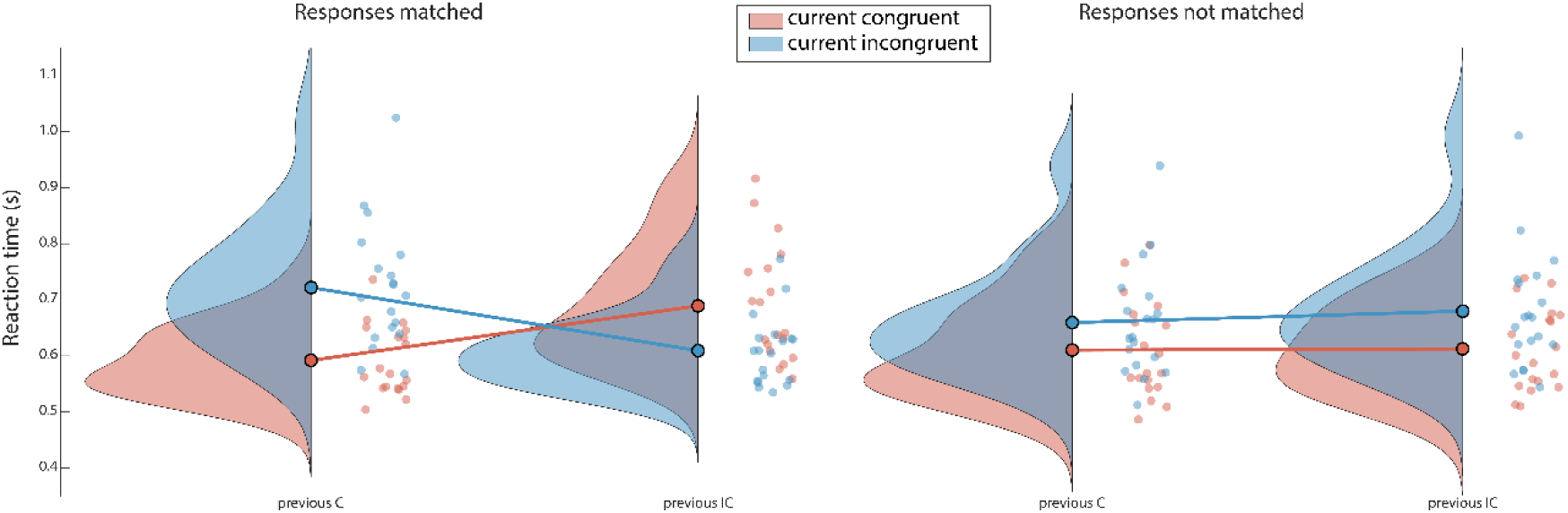
The Gratton effect is only present when two sequential trials require the same response. Graphs show probability density, box plots denote median, 1^st^ and 3^rd^ quartiles and 1.5 times the interquartile range (IQR). Dots correspond to data points of individual subjects.

**Figure S3.**
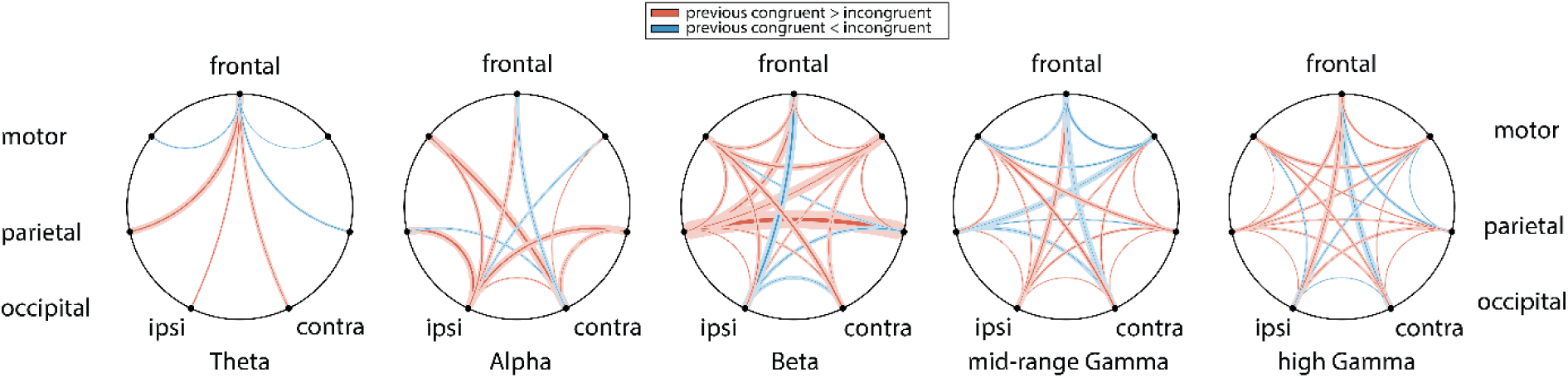
Coherence in the task-relevant network for all connections with p>0.05 (uncorrected). The relative difference in coherence for previous congruent versus previous incongruent trials in the pre-cue window. Solid lines indicate group average, with the thickness of the lines proportional to the ratio between coherence in congruent and incongruent trials; transparent lines indicate standard deviation. Red (blue) denotes larger (smaller) coherence for previous congruent versus incongruent. Ipsi: ipsilateral to cued response hand; contra: contralateral to cued response hand.

